# *In vivo* rat-brain mapping of multiple gray matter water populations using nonparametric D(*ω*)-*R*1-*R*2 distributions MRI

**DOI:** 10.1101/2024.06.07.597866

**Authors:** Maxime Yon, Omar Narvaez, Daniel Topgaard, Alejandra Sierra

**Author notes:** **Corresponding author details:** Dr. Maxime Yon.

## Abstract

Massively Multidimensional Diffusion MRI combines tensor-valued encoding, oscillating gradients, and diffusion-relaxation correlation to provide multicomponent sub-voxel parameters depicting the tissue microstructure. This method was successfully implemented *ex vivo* in micro-imaging systems and in clinical conditions but with a reduced diffusion frequency (*ω*) range due to the use of classical tensor-valued encoding. We demonstrate here its preclinical *in vivo* implementation with a protocol of 389 contrast images probing a wide diffusion frequency range of 18 to 92 Hz at *b*-values up to 2.1 ms/µm^2^ enabled by the use of modulated gradient waveforms and combined with multislice high-resolution and low-distortion EPI acquisition with segmented and full reversed phase-encode acquisition. This framework allows the identification of diffusion *ω*-dependence in the rat cerebellum and olfactory bulb gray matter (GM) and the parameter distributions are shown to resolve two water pools in GM with different diffusion coefficients, shapes, *ω*-dependence, relaxation rates, and spatial repartition whose attribution to specific microstructure could modify the current understanding of the origin of restriction in GM.

## Introduction

Magnetic resonance imaging (MRI) directly probes the biological tissue structure *in vivo* and non-invasively with spatial resolution in the order of hundreds of micrometers in clinical setup^1,2^ to tens of micrometers in preclinical conditions^3,4^. MRI contrasts, however, indirectly depict the tissue’s microstructure at a much thinner scale relying on various mechanisms such as water molecules self-diffusion or nuclear magnetic relaxation^5^. Diffusion^6^ MRI imparts contrast based on water molecule diffusive displacements modulated by the restriction and hindrance due to cell boundaries and intracellular organelles and is sensitive to a few micrometers scale microstructure^7–9^. Longitudinal and transversal relaxation rates depict variations in the local chemical water environment. They are sensitive to chemical and magnetization transfer with metabolites and macromolecules^10,11^, intercompartmental exchange^12^, and paramagnetic relaxation from iron or injected contrast agents^13^. Relaxation and diffusion contrast are complementary and can be combined with multidimensional MRI measuring jointly the multiple contrasts and allowing the determination of their correlation^14,15^. The analysis of such multidimensional data can allow for disentangling the signatures of different water pools within a single voxel, thus precisely characterizing the microstructure when heterogeneities are present below the voxel scale^15–17^.

Tensor-valued diffusion encoding^18^ can also disentangle water pools with different diffusion sizes, shapes, or orientations at a sub-voxel level.^19,20^ It allows expressing the MRI voxel as a sum of diffusion tensors (**D**) or diffusion tensor distribution (DTD)^21^ computed with either parametric^21–23^ or nonparametric approaches^24,25^. The nonparametric DTD approach was also enriched by incorporating longitudinal and transversal relaxation rates (*R*_1_ and *R*_2_)^26–28^ via variable repetition times (τ_R_) and/or echo times (τ_E_) and yielding to nonparametric **D**-*R*_1_-*R*_2_ distributions. DTD and DTD-based multidimensional diffusion found applications in clinical^19,28–32^ and preclinical^33,34^ MRI.

In parallel, tensor-valued encoding was enriched by frequency-dependent diffusion^35–37^ allowing the correlation between the diffusion shape and the diffusion frequency dependence sensitive to restriction. The diffusion frequency-dependence (*ω*)^38,39^ was then included in the multidimensional diffusion framework by transforming the diffusion encoding tensor (**b**)^40–43^ in a diffusion encoding spectrum **b**(*ω*)^20,30,37,44^ and yields frequency-dependent multidimensional diffusion **D**(*ω*)-*R*_1_-*R*_2_. On the MRI acquisition side, the non-zero *ω* content of the tensor-valued diffusion encoding due to the use of various gradient waveforms of different durations allows exploring the *ω-*dependence of **b**(*ω*) in a narrow frequency range. The seminal experiments were realized *ex vivo* in microimaging MRI with a high-gradient strength of 3 T/m and allowed to evidence *ω-*dependence in the various phantoms, rat brain-specific structures, and tumors^45,46^. The direct transfer of such an approach in clinical scanners with lower gradient strengths limits the frequency range and did not allow to observe restriction effects in the first two studies^47,48^. High-performance gradient hardware of micro-imaging systems allows for increasing the *ω*-span by relying on “double rotation” gradient waveforms^49^, inspired by both solid-state NMR^50,51^ and oscillating gradient spin echo (OGSE) MRI^35^. In current micro-imaging systems, these modulated gradient waveforms allow accessing diffusion frequencies up to 1 kHz at low *b*-values and up to 300 Hz at *b*-values over 4 ms/µm^2^ revealing important *ω*-dependence in specific mouse brain regions such as the cerebellar gray matter^47^.

The purpose of this article is to describe the implementation of massively multidimensional diffusion-relaxation correlation (MMD-MRI) *in vivo* in a preclinical setting with a moderated gradient strength of 760 mT/m with multislice acquisition and a high *ω*-span allowed by the use of modulated gradient waveforms on a rat brain. The various parameter maps produced by this framework will be detailed with a special focus on *ω*-dependent maps acquired for the first time *in vivo* with a wide frequency range of 18 and 92 Hz at *b*-values up to 2.1 ms/µm^2^. We will also illustrate here the specific signal distributions in white matter (WM), gray matter (GM), and cerebrospinal fluid (CSF). The GM distribution presenting two components will be further segmented to produce specific parameter maps. In the idea of open research, all the code used in this article is available on GitHub. The MRI sequence is available upon request. This article will also highlight the versatility and user-friendliness of the MRI sequence to promote its usage.

## Materials and methods

### Animal handling

All experiments were approved by the Animal Committee of the Provincial Government of Southern Finland following the guidelines established by the European Union Directives 2010/63/EU. The MRI acquisitions were performed on a healthy male rat Sprague-Dawley (Harlan Netherlands B.V.) aged 9 weeks and weighing 400 grams. Animals were housed in individual cages in a controlled environment (constant temperature 22 ± 1 °C, humidity 50–60%, lights on 07:00–19:00 h) with free access to food and water. The anesthesia was induced by ∼5% isoflurane and then kept with ∼2% isoflurane in a mix of 30/70% oxygen/nitrogen. The rodent respiration was monitored during the entire experiment and maintained at 35–60 breaths per minute by adjusting the isoflurane flux. The animal temperature was also monitored by a rectal sensor and maintained between 36 and 37 °C by a heated water pad placed under the animal.

### MRI acquisitions

All the acquisitions were performed on a horizontal Bruker PharmaScan® 7 Tesla preclinical scanner equipped with a 3-axis gradient system with a 90 mm inner diameter capable of delivering a gradient strength of 760 mT/m with a slew rate of 6840 T/m/s per axis and running Paravision 6.01. A ^1^H actively decoupled volume coil was used for RF transmission and combined with an anatomically shaped receive-only quadrature rat head surface coil (Bruker Biospin®). The rat heads were fixed with tooth and ear bars and the acquisitions were performed without respiratory triggering.

The acquisitions were performed using the Bruker Spin Echo-Echo Planar Imaging (SE-EPI) sequence customized with variable repetition time delay (τ_R+_), variable echo time delays (τ_E+_), and diffusion encoding with two identical self-refocusing gradient waveforms of various durations placed on each side of the refocusing pulse according to the sequence scheme presented in Fig. 1a. The EPI acquisitions were performed with a matrix size of 129×80×5, and a Field of View (FoV) of 29×20×5 mm^3^ leading to a spatial resolution of 250×250×700 µm^3^ with a 300 µm slice gap. A readout bandwidth of 35.7 kHz, the use of double-sampling^52^, two segments, and a phase-encoded partial Fourier factor of 1.4 led to a minimum τ_E_ without diffusion encoding of 21 ms and a bandwidth of 2.77 kHz in phase encode direction. Two repetitions were used to acquire reversed phase-encode blips images to perform top-up post-processing.

**Figure 1:**
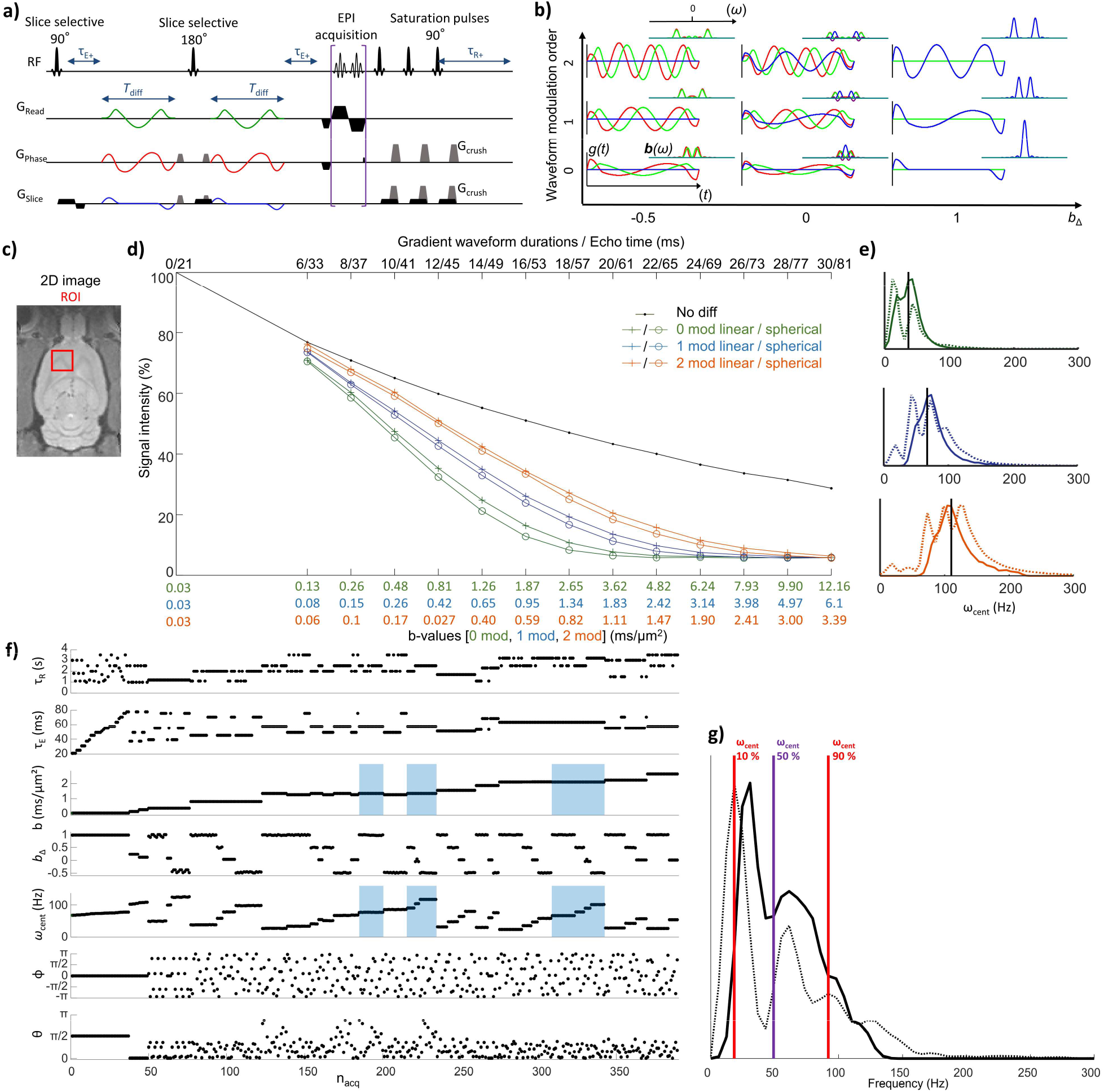
a) SE-EPI pulse sequence scheme with the integration of variable echo time delays (τ_E+_) and repetition time delay (τ_R+_), and tensor-valued diffusion encoding (τ_diff_). b) Gradient waveforms and corresponding encoding spectra **b**(*ω*) on the top right for *b*_Δ_ = -0.5, 0, 1 and modulated up to 2^th^ order. c) rat brain and selected ROI for signal integration. d) Signal decay curves in the ROI presented in panel c. In black, without diffusion weighting, in orange, blue, and green with diffusion weighting from the 0, 1, and 2 order waveforms of durations between 6 to 30 ms. Cross markers indicate linear encoding and circle markers spherical encoding. e) *ω* and *ω*_cent_ distributions in dashed and continuous lines respectively corresponding to the acquisition performed with the 0, 1, and 2 order waveforms. f) MMD protocol of 389 images, the blue rectangles highlight the images acquired with the 1st-order waveforms. g) Corresponding *ω* and *ω*_cent_ distributions in dashed and continuous lines respectively.

### Frequency-dependent tensor-valued diffusion encoding

The description of the encoding spectrum **b**(*ω*) requires the definition of a few mathematical notations which have been detailed in previous publications ^45,46,53^ and will only be briefly summarized in this article to allow self-sufficient reading.

The encoding spectrum **b**(*ω*) is calculated by the velocity autocorrelation function^54^ suitable when the narrow gradient pulse approximation is violated.^55^ It include both the gradient waveforms and the imaging gradients **g**(t) of duration τ via the time-dependent dephasing vector **q**(t) and its Fourier transform **q**(*ω*) according to:

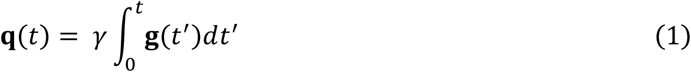

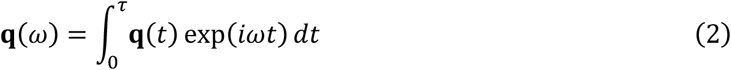

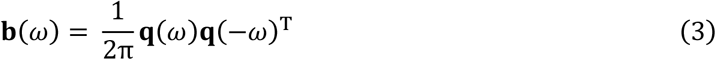

The encoding spectrum **b**(*ω*) is characterized by its *b*-value, its encoding anisotropy *b*_Δ_^41^, and its centroid frequency *ω*_cent_ ^56^ according to:

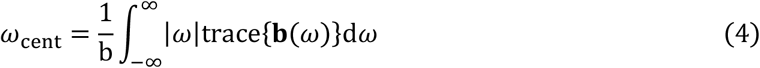

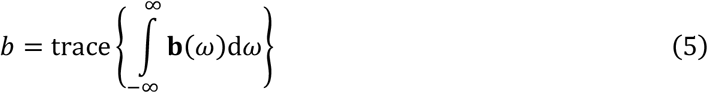

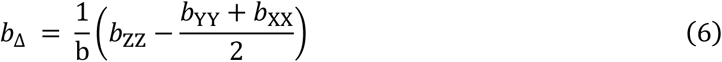

where *b*_XX_, *b*_YY_, and *b*_ZZ_ are the eigenvalues of the b-tensor ordered according to the convention |*b*_ZZ_– *b*/3| > |*b*_XX_–*b*/3| > |*b*_YY_–*b*/3|. The diffusion encoding was performed with two identical self-refocusing gradient waveforms placed on both sides of the refocusing pulse of the SE-EPI sequence as shown in Fig 1a. The waveforms used in this study are depicted in Fig 1b, due to the limited available gradient strength only two orders of modulation were used to allows reaching high *b*-values with reasonable diffusion waveforms duration^49^. In Fig 1b, the waveforms are normalized to a constant *b*-value per duration leading to constant integrals of the **b**(*ω*) spectra presented on the top right of each waveform.

### Massively multidimensional acquisition protocol

The acquisition protocol designed for this study is presented in Fig 1f. It is composed of 389 images with τ_R_ ranging from 1 to 3.5 s, τ_E_ from 21 to 77.5 ms, *b*-values from 0.03 to 2.65 ms/µm^2^, *b*_Δ_ of -0.5, 0, 0.5, *ω*_cent_ from 46 to 172 Hz for *b*-weighted images for an acquisition duration of 17 minutes. The total acquisition duration for the MMD dataset taking into account the two segments of the EPI acquisition and the full acquisition of blip-up and down images for the entire protocol led to an acquisition time of 1 hour and 9 minutes.

The table used to create such an acquisition protocol is presented in Table 1. The heuristic guidelines to design this protocol are presented in the result section.

**Table 1:**
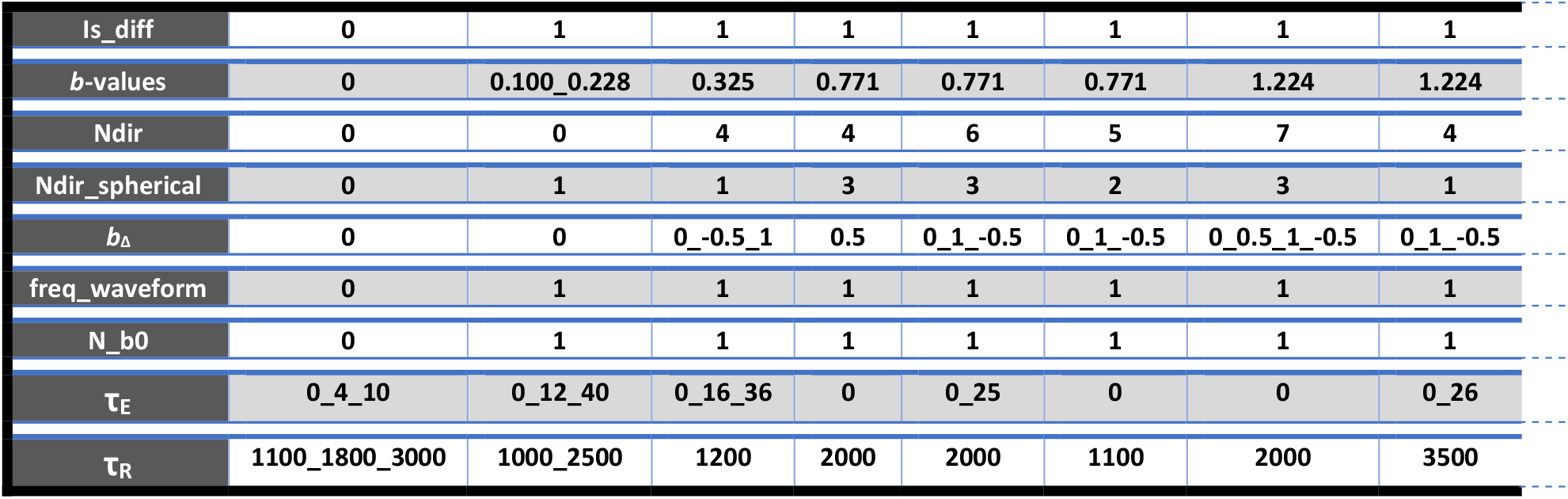
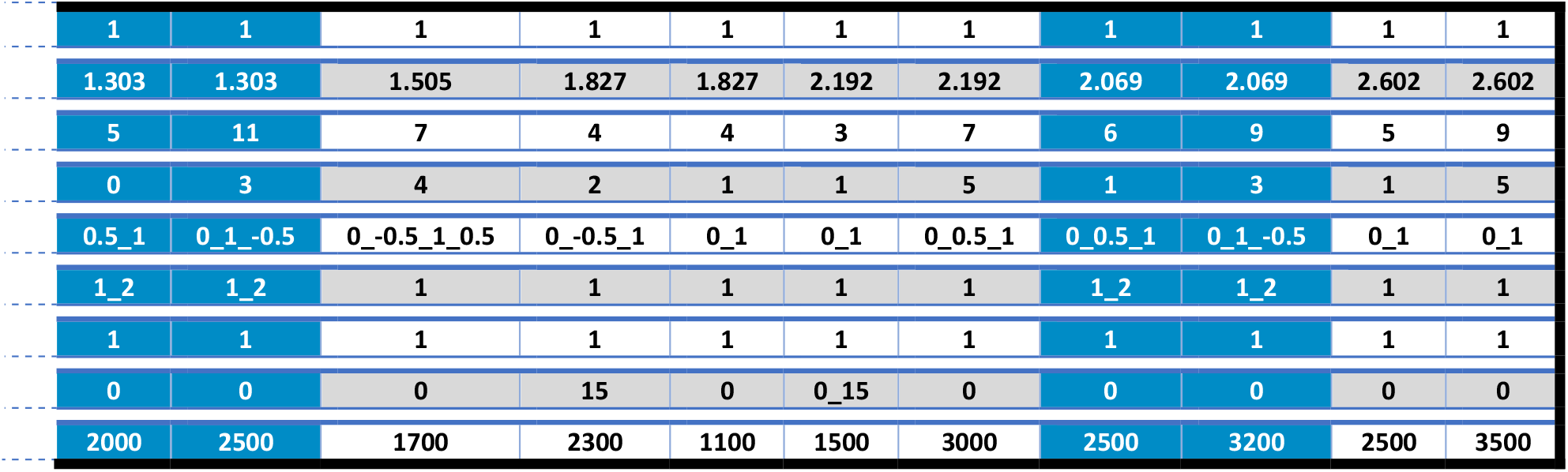
Table used for protocol generation. The blue columns contain the 1^st^ order waveform experiments.

|  |  |  |  |  |  |  |  |  |
| --- | --- | --- | --- | --- | --- | --- | --- | --- |
| $is\_diff$ | 0 | 1 | 1 | 1 | 1 | 1 | 1 | 1 |
| $b$-values | 0 | 0.100_0.228 | 0.325 | 0.771 | 0.771 | 0.771 | 1.224 | 1.224 |
| $N_{dir}$ | 0 | 0 | 4 | 4 | 6 | 5 | 7 | 4 |
| $N_{dir\_spherical}$ | 0 | 1 | 1 | 3 | 3 | 2 | 3 | 1 |
| $b_\Delta$ | 0 | 0 | 0_-0.5_1 | 0.5 | 0_1_-0.5 | 0_1_-0.5 | 0_0.5_1_-0.5 | 0_1_-0.5 |
| $freq\_waveform$ | 0 | 1 | 1 | 1 | 1 | 1 | 1 | 1 |
| $N_{b0}$ | 0 | 1 | 1 | 1 | 1 | 1 | 1 | 1 |
| $\tau_E$ | 0_4_10 | 0_12_40 | 0_16_36 | 0 | 0_25 | 0 | 0 | 0_26 |
| $\tau_R$ | 1100_1800_3000 | 1000_2500 | 1200 | 2000 | 2000 | 1100 | 2000 | 3500 |

|  |  |  |  |  |  |  |  |  |  |  |
| --- | --- | --- | --- | --- | --- | --- | --- | --- | --- | --- |
| 1 | 1 | 1 | 1 | 1 | 1 | 1 | 1 | 1 | 1 | 1 |
| 1.303 | 1.303 | 1.505 | 1.827 | 1.827 | 2.192 | 2.192 | 2.069 | 2.069 | 2.602 | 2.602 |
| 5 | 11 | 7 | 4 | 4 | 3 | 7 | 6 | 9 | 5 | 9 |
| 0 | 3 | 4 | 2 | 1 | 1 | 5 | 1 | 3 | 1 | 5 |
| 0.5_1 | 0_1_-0.5 | 0_-0.5_1_0.5 | 0_-0.5_1 | 0_1 | 0_1 | 0_0.5_1 | 0_0.5_1 | 0_1_-0.5 | 0_1 | 0_1 |
| 1_2 | 1_2 | 1 | 1 | 1 | 1 | 1 | 1_2 | 1_2 | 1 | 1 |
| 1 | 1 | 1 | 1 | 1 | 1 | 1 | 1 | 1 | 1 | 1 |
| 0 | 0 | 0 | 15 | 0 | 0_15 | 0 | 0 | 0 | 0 | 0 |
| 2000 | 2500 | 1700 | 2300 | 1100 | 1500 | 3000 | 2500 | 3200 | 2500 | 3500 |

### Post-acquisition preprocessing

The images were reconstructed in Paravision 6.01, then the full dataset was split into reversed phase-encode blips series. Each series was denoised with random matrix theory^57,58^ implemented in Python as well as Gibbs ringing removal^59^ implemented in MRTrix3^60^. Motion detection was then performed on Matlab® (Natick, Massachusetts: The MathWorks Inc.) with the “imregtform” function using the multimodal optimizer for translational motion only. The detected motion was then smoothed with a cubic smoothing splines function with a smoothing parameter of 10^−6^. The average of the reversed phase-encode blips curves was then used to apply identical translational motion correction on both image series. The first steps of the post-processing are illustrated in Fig 2a.

**Figure 2:**
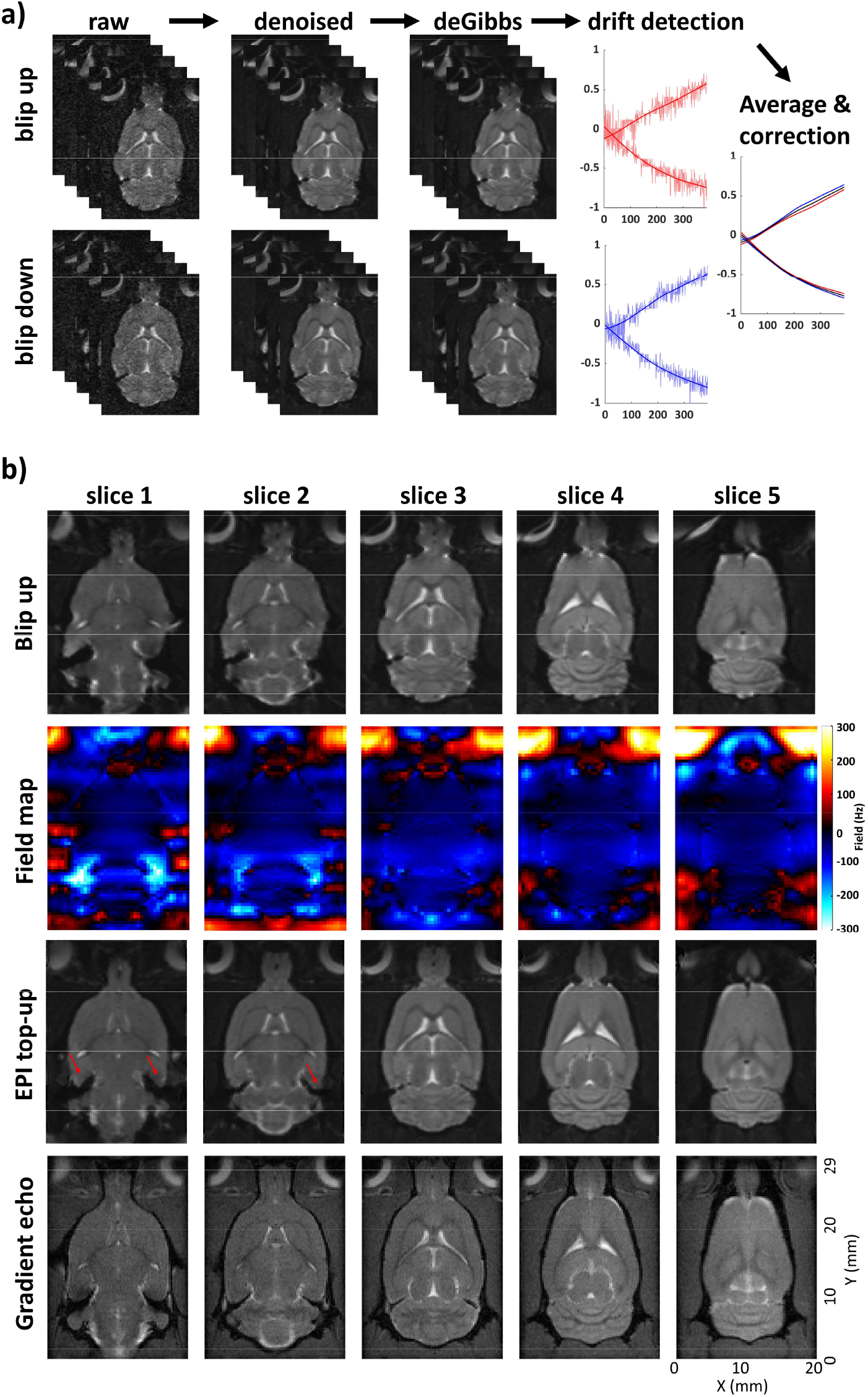
Post-acquisition preprocessing: a) of the reversed phase-encode blips series with, from left to right, random matrix theory denoising, Gibbs-ringing artifact removal, and translational motion detection, smoothing, averaging, and correction and b) from top to bottom: the blip up only images, the field maps calculated for every slice, the top-up corrected images and the corresponding slices imaged by a gradient echo sequence for comparison. The red arrows point to imperfectly corrected areas inducing signal drops and geometrical distortions.

Data was collected with reversed phase-encode blips, resulting in pairs of images with distortions going in opposite directions. From these pairs the susceptibility-induced off-resonance field was estimated using a method similar to that described in Andersson et al.^61^ as implemented in FSL^62^ and the two images were combined into a single corrected one. The result of this correction and its comparison with gradient echo image using the Fast Low Angle shot (FLASH) sequence^63^ is shown in Fig 2b.

### Multidimensional diffusion data inversion

The nonparametric Monte Carlo inversion of the multidimensional dataset was performed with Matlab using the *md-mri* Toolbox^64^ according to the following equation taking into account the encoding spectrum **b**(*ω*): ^45,46^

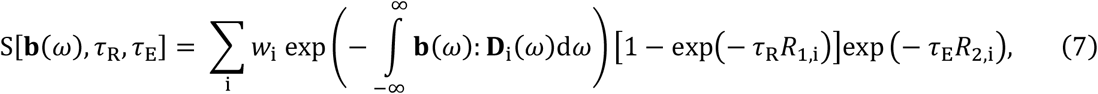

allowing to obtain the frequency-dependent self-diffusion tensor **D**(*ω*)^39^ for each of the components. The equation is made tractable by approximating **D**_i_ (*ω*) as axisymmetric tensors as:^46^

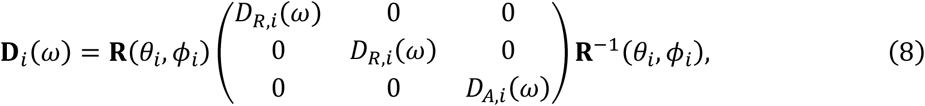

where **R**(*θ*_*i*_, *ϕ*_*i*_) is a rotation matrix and *ω*-the dependent radial and axial eigenvalues D_R,i_(*ω*) and D_A,i_(*ω*) are defined by:

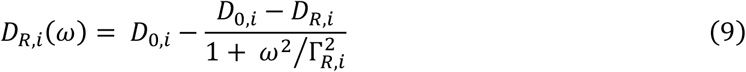

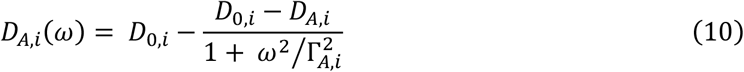

with *D*_0,i_ the diffusivity at high frequency, and Γ_A,i_ and Γ_R,i_ the rate of change of axial and radial diffusivities. The frequency-dependent isotropic diffusivity *D*_iso_(*ω*) and normalized diffusion anisotropy^65^ *D*_Δ_(*ω*) can be calculated by:

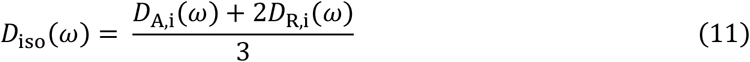

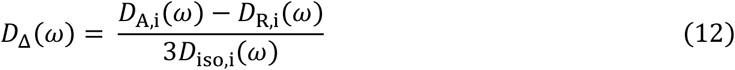

The inversion was performed with the limits: 5·10^−12^ m^2^s^−1^ < D_0/A/R_ < 5·10^−9^ m^2^s^−1^, 0.1 s^−1^< Γ_A/R_ < 10^5^ s^−1^, 0.1 s^−1^ < *R*_1_ < 4 s^−1^, and 4 s^−1^ < *R*_2_ < 100 s^−1^, 20 steps of proliferation, 20 steps of mutation/extinction, 200 input components per step of proliferation and mutation/extinction, 10 output components. The bootstrapping was performed by 100 repetitions using random sampling with replacement to compute the parameter maps and 1000 repetitions to obtain the ROIs distributions.

For generating parameter maps, the rich information in the ***D***(*ω*)-*R*_1_-*R*_2_-distributions is first collapsed to the mean values of the ensemble of solutions and then further condensed into median (E[x]), variances (V[x]), and covariances (C[x,y]) over relevant dimensions distributions^25^. The 2D *D*_iso_-*D*_Δ_^2^ space is divided into three bins to allow for image segmentation by coding the per-bin signal fractions *f*_bin1_, *f*_bin2_, and *f*_bin3_ into RGB color and extract the bin-specific diffusion metrics which in the brain are characteristic of white matter (WM), grey matter (GM) and cerebrospinal fluid (CSF)^33^. The bins boundaries were defined according to: bin1: *D*_iso_ < 2.5×10^−9^ m^2^s^−1^ and *D*_Δ_^2^ > 0.25; bin2: *D*_iso_< 2.5×10^−9^m^2^s^−1^ and *D*_Δ_^2^ < 0.25; and bin3: *D*_iso_>2.5×10^−9^m^2^s^−1^.

The frequency-dependent behavior of the isotropic diffusion *D*_iso_ and the shape parameter *D*_Δ_^2^ is quantified by the difference between the parameters calculated at high and low frequencies. Since the highest frequencies are achieved only at low *b*-value, the frequency range considered in the difference is restrained to the 10 and 90 percentiles of the *ω* values corresponding to 18 and 92 Hz and represented in Fig. 1g by the red lines. The frequency dependence parameters (Δ_*ω*/2π_) are calculated according to^45^:

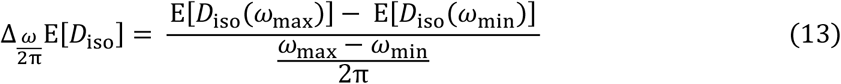

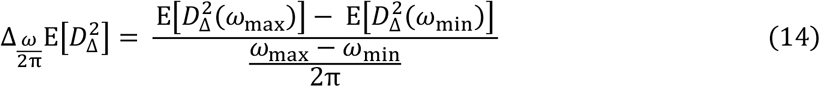

The correlation plots showing the distributions are obtained by projecting and mapping the weights of the discrete components onto 64×64 meshes in the corresponding 2D parameter space using 1×1 grid points Gaussian kernel.

## Results

The nonparametric Monte Carlo inversion algorithm used in this study allows for sparse sampling of the parameter space and thus complete freedom in the acquisition protocol design. However, in practice, hardware constraints, limited SNR, and compromise between the sampling of different parameters limit the parameter ranges that can be probed. This is especially true for the sampling of high diffusion frequencies and high *b*-values restraining the use of highly modulated gradient waveforms with limited gradient strengths and short transversal relaxation time (T_2_) samples. This compromise is depicted in Fig 1d showing the signal decay curves obtained with gradient waveforms modulated up to the second order and recorded *in vivo* in the rat brain ROI presented in Fig 1c. The signal is normalized to 100 for the shortest echo time (τ_E_) of 21 ms allowed by the SE-EPI sequence without diffusion encoding. The black curve presents the T_2_ decay from 21 to 81 ms τ_E_ without diffusion weighting and the orange, blue, and green curves present the signal decay for the 0, 1, and 2 modulation waveforms respectively presented in Fig 1b. The waveforms’ duration is 6 to 30 ms and utilizes 99.9 % of the maximum available gradient strength. For each modulation order, two curves corresponding to linear and spherical encodings (bΔ of 0 and 1) are displayed. The vertical distance between the black and colored curves shows the diffusion weighting obtained for each of the waveforms which decreases with the increase of modulation order. The distance between two curves of the same color shows the difference between linear and spherical encoding for the defined ROI containing both anisotropic and isotropic diffusion water population. This gap is proportional to the diffusion weighting. Fig 1e compares the *ω* and *ω*_cent_ distributions for each of the three modulated waveforms and for durations between 6 to 30 ms. The *ω*-distribution presented in the dashed line corresponds to the sum of all b(*ω*) obtained by calculating the trace of the encoding spectra **b**(*ω*). A width of the *ω* distribution matching the one of the *ω*_cent_ distribution means that the frequency dispersion is mainly due to the variation of *ω*_cent_ and that the protocol is specific to the frequency changes. However, if the *ω* distribution width is dominating, the acquisition is thus not very frequency-specific^47^. The black lines represent the center of the *ω*_cent_ distribution whose values are 37, 67, and 110 Hz for the waveform modulation orders 0, 1, and 2 respectively.

These data allow guiding the creation of a sampling protocol compromising between high diffusion weighting and high-frequency sampling via the use of both the 0 and 1^st^ order waveform modulations presented in Fig 1f and including 389 images for an acquisition duration of 17 minutes. Fig 1f presents the experimental values calculated from the acquired data and considers all the imaging gradients and sequence delays including the EPI acquisition up to the k-space center. Blue rectangles highlight the use of the 1st-order gradient waveforms in Fig 1f allowing higher diffusion frequencies up to 116 Hz for moderate *b*-values of 1.3 ms/µm^2^ and 100 Hz for *b*-values of 2.1 ms/µm^2^ compared to the 0-order waveforms. However, these high frequencies at high *b*-values are also correlated with long waveform durations up to 21 ms and thus long τ_E_. The use of the 0^th^ order waveforms for the acquisition of the maximum *b*-value of 2.65 ms/µm^2^ allows to reduce the waveform duration to 18 ms and thus the τ_E_ by twice the difference: 6 ms. To avoid overfitting in the *ω*-dependent diffusion parameters estimation, the frequency limits are constrained to the 10 to 90 % of frequency sampling density corresponding to 18 to 92 Hz shown in Fig 1g by the red lines. The protocol was created with the xls table presented in Table 1 and interpreted by a Matlab code whose convention will be explained here to show the versatility of the sequence design and share the protocol in a convenient and compact form. Each column of the table defines a set of experiments, and the full protocol is defined by the addition of all the columns.

Each line of the table will be detailed in the following list:

– 1, is_diff: allows to perform acquisitions without diffusion weighting when set to 0 and thus record images at minimum τ_E_ which was shown to decrease the estimation bias of the Monte Carlo inversion^66^.
– 2, *b*-values: contain the targeted *b*-value(s) without considering the imaging gradients. The waveform duration will be set to the smallest value rounded to the millisecond allowing it to reach the targeted *b*-value. If the cell contains multiple *b*-values separated by a “_” character as in the second column the waveform duration will be set to the value allowing to reach the highest *b*-value.
– 3, Ndir: contains the number of diffusion directions for all *b*_Δ_ values different from 0.
– 4, Ndir_spherical: contains the number of diffusion directions for spherical encoding (*b*_Δ_ = 0). This distinction allowed a reduced number of spherical encoding directions since this encoding probes the diffusion in three orthogonal directions each time.
– 5, *b*_Δ_: encoding anisotropy *b*_Δ_ value, all values between -0.5 and 1 are accepted.
– 6, freq_waveform: modulation order of the waveform, comprised between 0 to 5. When multiple values are specified, the waveforms are normalized to allow identical diffusion encoding durations.
– 7, N_b0: number of b0 image for the column
– 8, τ_E_: additional τ_E_ in milliseconds which is equally split on each side of the spin echo outside of the diffusion encoding in τ_E+_ delays shown in Fig 1a.
– 9, τ_R_: Targeted repetition time (τ_R_) corresponding to the τ_R+_ delay shown in Fig 1a.

To limit the gradient heating and avoid acquisition bias, the acquisition is performed in randomized order and the table order might not exactly correspond to the order of Fig 1f in which the protocol is sorted by *b*-values, *ω*_cent_ values, and *b*_Δ_ values. In multislice acquisition, the minimum τ_R_ delay limits the number of slices since the acquisition of the additional slices needs to fit into the τ_R+_ delay. In this study, the minimum τ_R_ of 1 second allows the acquisition of 5 slices.

Fig 2a presents the post-processing steps applied to each reversed phase-encode blips image series starting with random matrix theory denoising followed by Gibbs-ringing artifact removal and translational motion correction. The motion curves show that smooth translational motion dominates the detected motion and such behavior is consistent with motion induced by field drift probably due to gradient heating. Fig 2b presents the SE-EPI images before the top-up correction, the obtained magnetic field maps, and the top-up corrected images as well as gradients echo reference images. The top-up correction based on the full acquisition of the reversed phase-encode blips series is proven very effective with high-quality corrected images even if an unperfect image combination remains when the field deviation reaches more than 300 Hz and lead to signal drops and geometrical distortions indicated by the red arrows.

Fig 3 exhibits the parameter maps extracted from the ***D***(*ω*)-*R*_1_-*R*_2_-distributions corresponding to slice 4 of the *in vivo* rat brain. It includes the *S*_0_ and fractions maps, the per-voxel means E[x], variances V[x], and covariances C[x,y] of the *D*_iso_, *D*_Δ_^2^, *R*_1_, *R*_2_ and the rate of change with frequencies (Δ_ω/2π_) of *D*_iso_ and *D*_Δ_^2^ as well as their corresponding bin resolved fractions. The definition of the parameters was already detailed in previous works^28,32,46^ and will only be briefly reminded here. Fig. 3a includes the *S*_0_ parameter maps corresponding to the signal intensity calculated by eq. 7 at infinite τ_R_, zero τ_E_, and zeros diffusion weighting, it is obtained by the sum of the *w*_i_ coefficients. The *S*_0_ map presents little contrast and only the CSF can be clearly distinguished even if GM seems in general a little brighter than WM. The fraction maps present the division of the 2D *D*_iso_-*D*_Δ_^2^ solution space into three bins whose boundaries are represented in Fig 3a and are detailed in the multidimensional diffusion data inversion section. This map allows to separate the WM, GM, and CSF into red, green, and blue fractions with the intermediate color representing composite voxels of WM + GM (yellow), WM + CSF (purple), or GM + CSF (turquoise)^29,33^. Fig 3b encompasses the means E[*D*_iso_], which correspond to the MMD estimate of conventional mean diffusivity^67^, E[*D*_Δ_^2^] is analogous to earlier metrics quantifying microscopic diffusion anisotropy^20,40,68–70^ and *R*_1_ and *R*_2_ correspond to longitudinal and transversal relaxation rates according to *R*_1_ = 1/T_1_ and *R*_2_ = 1/T_2_. The metrics *D*_iso_ and *D*_Δ_^2^ are *ω*-dependent according to eq. 11 and 12 respectively and are calculated for the low frequency of *ω* = 18 Hz. At this frequency, the E[*D*_iso_] map highlights the CSF while the E[*D*_Δ_^2^] map allows identifying the WM areas. The *R*_1_ and *R*_2_ maps provide additional contrast with increasing values between WM, GM, and CSF. The directionally-encoded color maps are obtained from the lab-frame diagonal values [*D*_*xx*_,*D*_*yy*_,*D*_*zz*_] normalized by the maximum eigenvalue *D*_33_. The rates of change with frequency (Δ_ω/2π_) are depicted in Fig 3c as maps for both *D*_iso_ and *D*_Δ_^2^ between the densely sampled frequencies of 18 and 92 Hz and calculated according to equations 11 and 12. The Δ_ω/2π_E[*D*_iso_] corresponds to the earlier Δ_*f*_ADC metric classically used in OGSE experiments^71^. The cerebellar granule cell layer and cerebellar molecular layer, as well as the olfactory bulb, are all highlighted by Δ_ω/2π_E[*D*_iso_] and Δ_ω/2π_E[*D*_Δ_^2^] with positive and negative values respectively indicating an increase of *D*_iso_ and a decrease in *D*_Δ_^2^ of with the increase of diffusion frequency which suggestive of restriction behaviors^47,71^. Outside of these regions, the Δ_ω/2π_E[*D*_iso_] also present low positive values in GM and negative values in the center of the lateral ventricles (lv) containing CSF while WM areas are characterized by zero values. The negative Δ_ω/2π_E[*D*_iso_] values are characteristic of the presence of coherent flow^48^. For each of those parameters the variance V[x] and variance C[x,y] maps shown in lines two and three report on various aspects of intravoxel heterogeneity and highlight voxels comprising multiple water populations with different diffusion and/or relaxation properties. High values of variance are found in the V[*D*_iso_] map in voxels containing a mixture of CSF with high diffusivity and WM or GM with a much lower *D*_iso_ value. The V[*D*_Δ_^2^] map is more sensitive to voxel with partial volume of highly anisotropic WM and more isotropic CSF of GM. The covariance maps highlight voxels with non-zero variances and correlated parameter values such as voxels containing partial volumes of both WM (low *D*_iso_ and high *D*_Δ_^2^) and CSF (high *D*_iso_ and low *D*_Δ_^2^)^28^.

**Figure 3:**
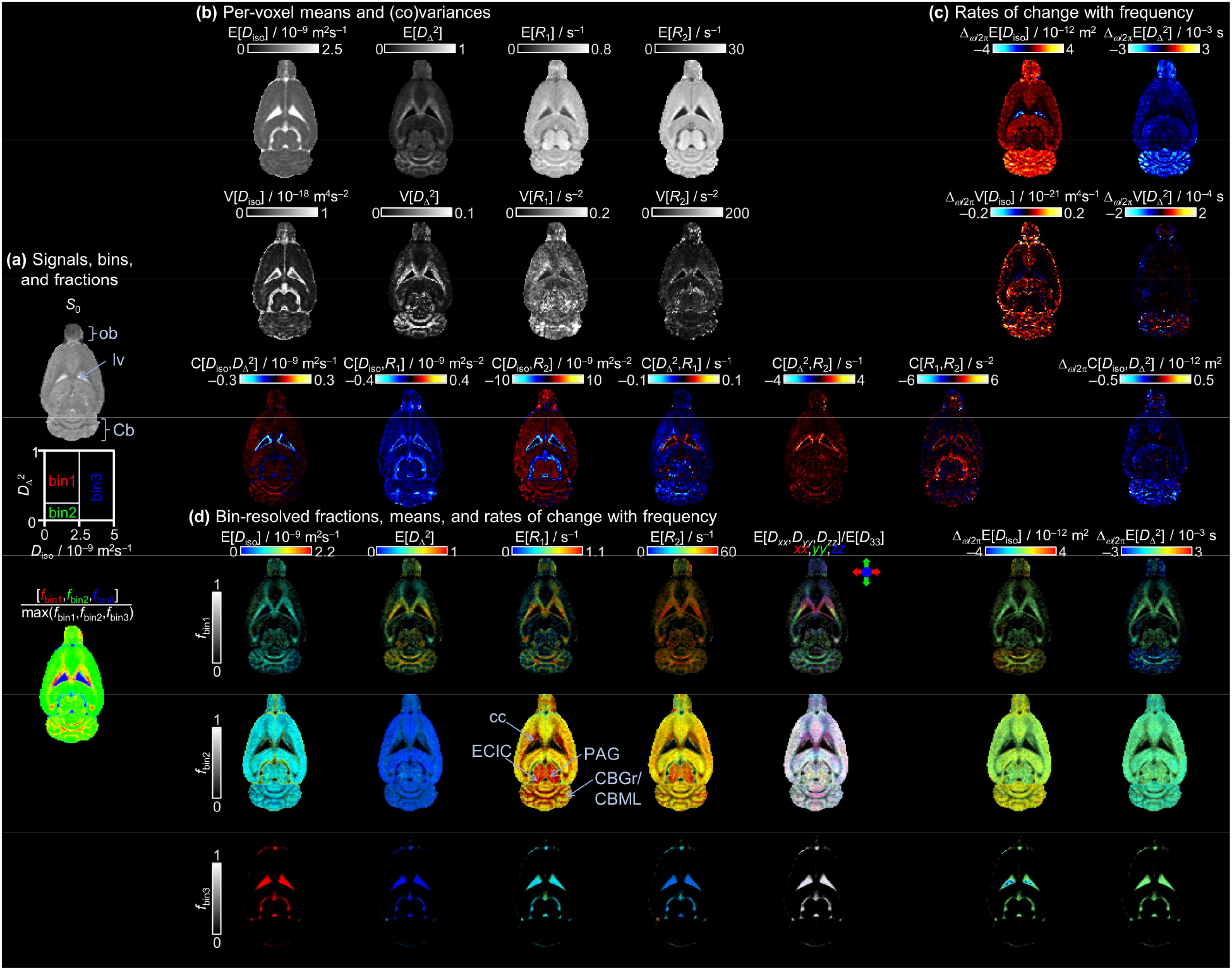
Parameter maps derived from the per-voxel ***D***(*ω*)-*R*_1_-*R*_2_-distributions of the *in vivo* rat brain slice 4. (a) Non-weighted signal *S*_0_ = *S*(*b*=0, *τ*_R_ → ∞, *τ*_E_ = 0) obtained by Eq. 7; graphical representation of the 1/2/3 bins in the 2D *D*_iso_-*D*_Δ_ ^2^ space calculated according to the bin limit defined in the multidimensional diffusion data inversion section; and bin-resolved signal fractions *f*_bin1/bin2/bin3_ coded into RGB color. The bin limits are selected to resolve white matter (WM), gray matter (GM), and cerebrospinal ?uid (CSF) at the low *ω* value of 18 Hz. Primary colors indicate voxels containing pure WM, GM, or CSF while mixed colors show voxels with partial volumes of WM + GM (yellow), WM + CSF (purple), or GM + CSF (turquoise). (b) Per-voxel means E[x], variances V[x], and covariances C[x,y] of the *D*_iso_, *D*_Δ_ ^2^, *R*_1_, and *R*_2_ dimensions. (c) Per-voxel rates of change with frequency (Δ_ω/2π_) means Δ_ω/2π_E[x], variances Δ_ω/2π_V[x], and covariances Δ_ω/2π_C[x,y] of the *D*_iso_, *D*_Δ_ ^2^ dimensions. The ω-dependent parameters (*D*_iso_ and *D*_Δ_ ^2^) were calculated at the low sampling frequency of *ω* = 18 Hz and their corresponding Δ_ω/2π_ values between *ω* = 18 and 92 Hz. (d) Bin-resolved signal fractions *f*_bin1/bin2/bin3_, means (E[x] and Δ_ω/2π_E[x]) coded into image brightness and color, respectively. The directionally-encoded color is based on the lab-frame diagonal values *D*_xx_, *D*_yy_, and *D*_zz_ normalized by the maximum eigenvalue *D*_33_. Specific brain regions such as the olfactory bulb (ob), the lateral ventricle (lv), the cerebellum (Cb), the corpus callosum (cc), the cingulum (cg), the periaqueductal gray (PAG), the external cortex of the inferior colliculus (ECIC), the cerebellar granule cell layer (CBGr) and cerebellar molecular layer (CBML).

The bin-resolved mean parameter values presented in Fig 3d report on the mean values associated with bin 1, bin 2, and bin 3 from top to bottom according to the three bins defined in Fig 3a.

Each map combines two orthogonal scales: the brightness intensity shows the relative signal fraction, and the color scale represents the mean value of the parameter. In the bin resolved E[*D*_iso_] maps, we observe low values below 1.5×10^−9^ m^2^/s (blue to green) for the bins associated with WM and GM while bin 3 corresponding to CSF shows high (red) values around 3×10^−9^ m^2^/s. In the bin resolved E[*D*_Δ_^2^] the highest values, above 0.3 are reported in the WM bin map, while the GM and CSG bin maps present only low values below 0.2 corresponding to isotropic diffusion. The per-bin relaxation rate maps show interesting contrast highlighting by high values the periaqueductal gray, the external cortex of the inferior colliculus, the cerebellar granule cell layer as well as the cerebellar and corpus callosum white matter areas. Supplementary Figure 5 shows the per-voxels and bins distributions of the parameter values over the entire brain. The last two columns of Fig 3d show the bin-resolved values of Δ_ω/2π_E[*D*_iso_] and Δ_ω/2π_E[*D*_Δ_^2^] respectively. High values of Δ_ω/2π_E[*D*_iso_] characterized by orange to red voxels indicating values above 1.5 ×10^−12^ m^2^ are primarily noted in the cerebellar gray matter both in the bin 1 and bin 2 maps and to a lesser extent in the olfactory bulb. The GM in bin 2 exhibits low values characterized by a mix of green to yellow colors (<1.5 ×10^−12^ m^2^) while WM in bin 1 is colored in pure green indicating values close to zero. The CSF bin in the lateral ventricles contains a few voxels of negative values of Δ_ω/2π_E[*D*_iso_] characteristic of coherent flow^48^. The Δ_ω/2π_E[*D*_Δ_^2^] map highlights mainly the bin 1 map with low negative values under -2 ×10^−3^ s in the cerebellar granule cell layer and olfactory bulb while only values of around -1 ×10^−3^ s colored in light blue are reported in the GM bin maps of the same regions.

This difference is explained by the low maximum value of *D*_Δ_^2^ (0.25) in bin 2 which does not allow much decrease with frequency increase.

Fig 4 presents the signal response and distributions corresponding to four ROIs located in different brain tissue regions as well as *S*_0_ maps corresponding to specific signal fractions. Fig 4a re-uses the fraction map already presented in Fig 3a to localize the 4 ROIs in WM, GM in the cortex, GM in the cerebellum, and CSF. The size of the ROIs is comprised of 1 to 20 voxels depending on the region homogeneity and the possibility of finding neighboring equivalent voxels. Fig 4b shows the experimental signal intensity in the four ROIs in black circles and the corresponding fit in colored dots back-calculated from the distributions and shows the good agreement between experimental points and fit as well as the overall decay behavior of each tissue. Fig 4c displays the corresponding ***D***(*ω*)-*R*_1_-*R*_2_-distributions in the *D*_iso_-*D*_Δ_^2^ plane, the *D*_iso_-*R*_1_ plane, and the *D*_iso_-*R*_2_ plane computed for five *ω*-frequencies equally spaced between 18 to 92 Hz with whiter shades of color with the increase of frequency. The first column depicts the distributions obtained in a voxel localized in the corpus callosum. In the *D*_iso_-*D*_Δ_^2^ plane, two contributions with *D*_Δ_^2^ values of 1 and 0.25 are visible, the one at *D*_Δ_^2^ = 1 is more intense and shifts the median value of the distribution to the high anisotropy fraction color-coded in red in the fraction map. The *D*_Δ_^2^ = 0.25 component could be due to spurious planar component artificially created by the inversion algorithm in short protocol and low SNR data^27,33^. The contour lines of different shades of red are well superimposed in both the *D*_iso_ and *D*_Δ_^2^ dimensions confirming the absence of *ω*-dependent behavior between 18 to 92 Hz in WM as already seen in the Δ_ω/2π_ parameter maps presented in Fig 3c. The second and third columns of Fig 4c show the distributions obtained in GM of the cortex and cerebellum respectively. The two distributions present two different contributions which can be roughly separated by a threshold at 0.5 × 10^−9^ m^2^/s in the *D*_iso_ dimension represented by the light blue line into two water pools: one with a low *D*_iso_ value (GM1) and one with a high *D*_iso_ value (GM2). The GM1 pool is characterized by lower isotropic diffusivity, slightly lower anisotropy, no *ω*-dependence, and higher *R*_1_ and lower *R*_2_ compared to GM2. The difference in *ω*-dependence is especially visible in the cerebellum distributions (third column) where the *ω*-dependence is more important than in the corpus callosum. The *ω*-dependence of the *D*_Δ_^2^ parameter is more difficult to assess in Fig 4c but seems to also show a reduced dependence of GM1 compared to GM2. The comparison of the 1D distributions of GM1 and GM2 over the entire brain is presented in Supplementary Fig 6 and shows more clearly this effect. While the correlation with *D*_iso_ allows identifying the *R*_1_ and *R*_2_ components belonging to each water pool, such identification based on the relaxation time alone would have been difficult due to their important overlap as shown in supplementary Fig. 6. The last column shows the distributions obtained in a CSF voxel characterized by a high *D*_iso_ value of around 3 m^2^/s and no anisotropy as well as a low *R*_2_ of around 4 s^-1^. Fig 4d and e show the percentage of *S*_0_ signal corresponding to GM1 and GM2 respectively. The GM1 fraction has a relative weight inferior to the one of GM2 with a maximum signal intensity of up to 30 % in the cerebellum and olfactory bulb and around 10 to 20 % in the rest of the brain GM. Focusing on the cerebellum the signal drops due to WM voxels are narrower in the GM1 map compared to the GM2 maps indicating an increased presence of the GM1 fraction in the granular layer. The supplementary figure 7 presents the normalized overlay of GM1 and GM2 fractions and allows visualization of these subtle differences.

**Figure 4:**
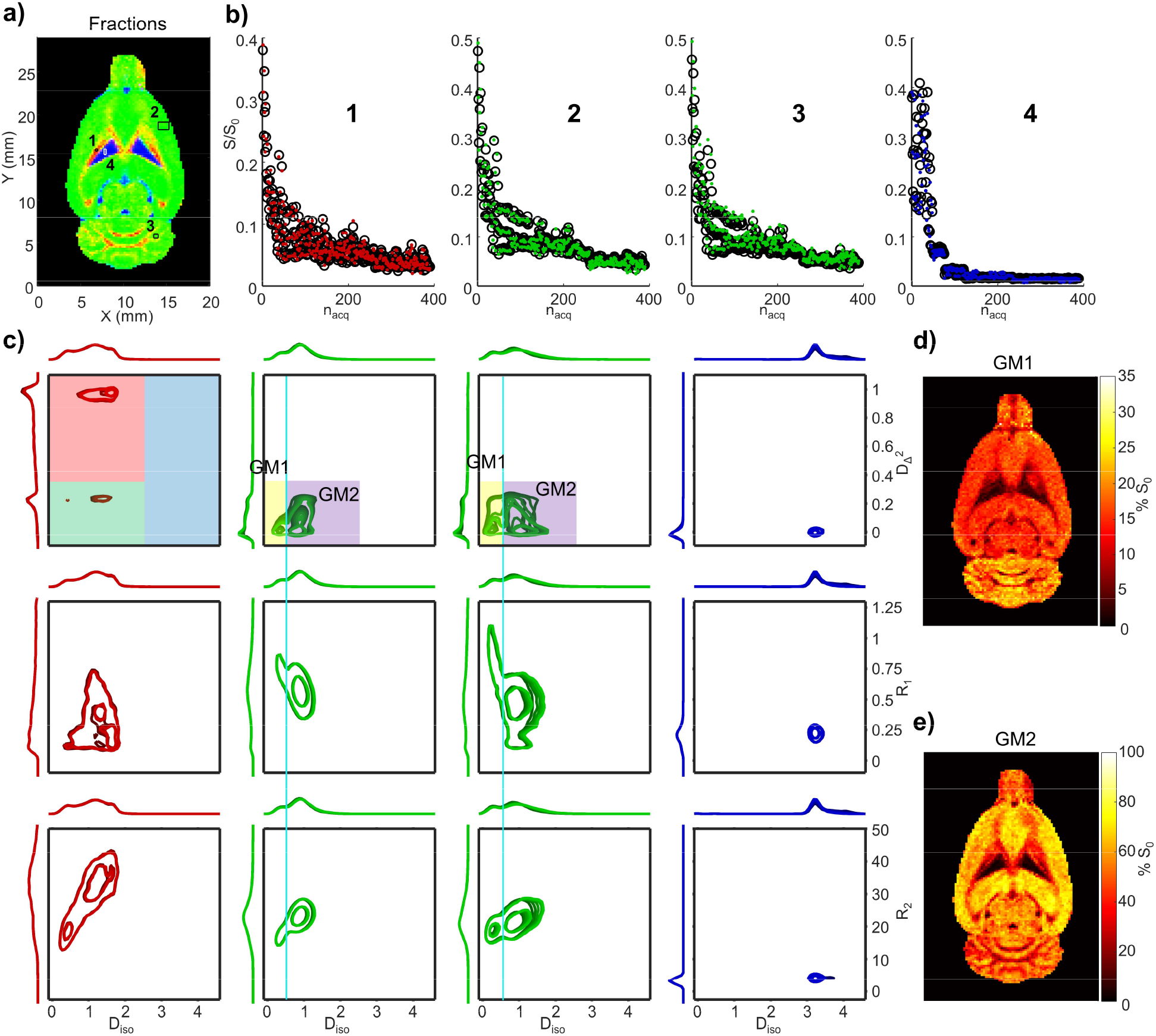
Experimental signal, fits, distributions, and additional binning maps. a) Fraction map indicating the positions of the ROIs: 1: in WM (single voxel), 2: in cortex GM (20 voxels), 3: in the cerebellum GM (4 voxels), and 4: in the CSF of the lateral ventricle (3 voxels). b) Experimental signal and median fit associated with the ROIs sorted to correspond to the protocol presented in Fig 1f. c) 2D distributions of each ROI with, from top to bottom the *D*_iso_-*D*_Δ_^2^ plane, the *D*_iso_-*R*_1_ plane, and the *D*_iso_-*R*_2_ plane computed at frequencies of 18, 36.5, 55, 73.5, and 92 Hz with whiter shades of color with the increase of frequency. The color of each of the 2D distribution lines is determined by the bin of the median value: red for bin 1, green for bin 2, and blue for bin 3. The bin boundaries are reminded in the top left 2D graph. An additional binning is displayed in the *D*_iso_-*D*_Δ_^2^ distribution of GM (columns 2 and three) with yellow and purple rectangles identifying two GM water pools GM1 and GM2 respectively. d) and e) present the corresponding maps of the fraction of the *S*_0_ signal corresponding to GM1 and GM2 respectively.

## Discussion

This article described the framework and guidelines allowing the acquisition of *in vivo* rat brains multislice MMD-MRI datasets with a wide diffusion frequency span allowed by the use of modulated gradient waveforms. The need for a compromise between the diffusion frequency span and T2-weighting controlled by the gradient waveform’s modulation order is especially emphasized. The protocol design is proven to be versatile and allows the use of various waveform modulation orders with or without normalization to allow the recording of the same b-weighting at various frequencies or maximal b-weighting with short τ_E_ with low-order waveforms. The acquisition framework was made user-friendly by the definition of the entire protocol based on a simple table as the one presented in Table 1 to allow the reader to create tailored acquisition schemes with either full MMD-MRI acquisition or a subset of it by keeping some of the parameter constants.

MMD-MRI is also combined with state-of-the-art ultrafast EPI-based acquisition and processing. The beneficial effect on image quality of top-up processing with full reversed phase-encode blips image acquisition is evidenced in all regions where the magnetic field inhomogeneities are inferior to 250 Hz and the processing code is made publicly available to increase its use which is still today not the default in the preclinical community^72^ partly due to the difficulty to link the preclinical images and parameters to the processing software designed for clinical data. Eddy-current correction is not part of the processing pipeline because no characteristic artifacts such as image distortions or rotation were noted, and also because the MMD-MRI does not use a classical shell encoding and the low number of images with identical repetition and echo times as well as diffusion waveform and duration preclude the use of the well-established algorithms based on opposite diffusion gradients acquisitions^73^. Note that in human MMD-MRI acquisitions, image registration-based algorithms are used for Eddy-current correction^29,48^. The only detected artifacts were smooth translations over time which are shown in Fig 2a and are probably due to magnetic field drift due to gradient heating. An image-based registration with a motion smoothing step was added to the preprocessing pipeline to correct it robustly. Overall, these developments as well as the open-access processing code are expected to increase the use of in and *ex vivo* MMD-MRI acquisitions.

Fig 3 to 4 evidence the wealth of information offered by the MMD-MRI frameworks providing access to nonparametric multicomponent **D**(*ω*)-*R*_1_-*R*_2_ distributions describing tissue microstructure at a sub-voxel level. Such a high-dimensionality framework was originally implemented *ex vivo* on a micro-imaging magnet but without using modulated gradient waveform and relying only on high gradient strength (3 T/m) to achieve diffusion frequencies from 50 to 150 Hz. Later on, the introduction of modulated gradient waveforms^49^ allowed to reach frequencies up to several hundred kHz^47^ with the same micro-imaging gradient system. The framework was also implemented and tested in clinical systems on human brains with a narrower frequency range of 6 to 21 Hz^47,48^ due to the limited gradient strength and the use of non-modulated gradient waveforms. Here the use of variable duration modulated gradient waveforms allows us to densely sample diffusion frequencies from 18 to 92 Hz with gradient waveform duration between 6 to 21 ms and a gradient strength of only 760 mT/m easily available in preclinical MRI. Diffusion frequency dependence suggestive of restriction was evidenced in the Δ_ω/2π_ parameter maps in both the cerebellum and olfactory bulb gray matter in agreement with previous *ex vivo*^45,71^ and *in vivo*^74,75^ studies in rat brain. The cerebellum GM is composed of three layers ordered from deep to superficial as follows: the cerebellar granule cell layer, the single layer of Purkinje cells, and the cerebellar molecular layer. The mean thickness of the cerebellar granule cell layer and cerebellar molecular layer in adult rats are around 150 µm and 220 µm respectively^76^. The Purkinje cell bodies are arranged as a unicellular discontinuous layer which is not expected to have a significant impact on the diffusion-weighted signal^75^. Previous studies reported higher restriction in the cerebellar granule cell layer compared to the cerebellar molecular layer, the Δ_ω/2π_E[*D*_iso_] and Δ_ω/2π_E[*D*_Δ_^2^] maps do not allow to distinguish them clearly. This could be due to the lowest frequency range of 18 to 92 Hz used in this study compared to frequencies up to 200 Hz in OGSE studies^71,74^. The restricted behavior was attributed to the granule cells whose mean diameter is approximately 7 to 10 µm^77^ and possess a thin cytoplasm which could lead to restriction in the frequency domain accessible in preclinical MRI. Indeed, in a rough approximation, neglecting size and shape distributions and exchange between water pools, a model of spherical confinement^78^ to a liquid can be applied with a diffusivity of 3 × 10^−9^ m^2^/s (corresponding to CSF diffusivity in Fig 4), a frequency range of 18 to 92 Hz and a threshold of observable variation of 20 % yield to a restriction sensitivity to a diameter range of 3 to 20 µm with a maximum at 7 µm.

Figure 4 also allows us to identify two water pools (GM1 and GM2) in GM both in the cortex and cerebellum with different diffusion and relaxation properties. The non-monoexponential decay induced by diffusion was already observed in GM^79–81^ but the assignment of each component to a specific microstructure remains difficult ^81^. Here, the full correlation between the parameters as well as the quantitative mapping over many brain regions provides a unique opportunity for a tentative assignment.

The GM2 fraction showed an important *ω*-dependence in the 18 to 92 Hz frequency range associated with restriction at 3 to 20 micrometers scale. The very small *ω*-dependence of the GM1 fraction can be associated with the absence of restriction effect or restriction at a scale outside of the sensitive range of the current measurement and thus compatible with a water pool restricted by cellular structures whose size is inferior to 3 µm. The low apparent diffusion coefficient of the GM1 fraction can be explained by either a low-viscosity water pool or the restriction by small structures. The differences in relaxation times of the two gray matter fractions remain more difficult to assign to specific microstructures. Indeed, while many studies show multicomponent T_2_ and sometimes T_1_ in white matter^82^, gray matter relaxation times are often described as a single component^83–85^. Note that without the correlation with the diffusion properties, the T_1_ and T_2_ distributions of gray matter presented in Supp Fig 6 would have led us to the same conclusion. However, in biological tissues T_1_ is well-known to be mainly dependent on the concentration of hydrated macromolecules^86,87^ and the lower T_1_ value of the GM1 fraction could thus indicate a higher concentration of macromolecules in this fraction. A comparison of the T_2_ values is more complex and no conclusion will be drawn from the T_2_ differences between the two fractions. The relative quantification and spatial distribution of gray matter fractions can also be taken into account as additional information to correlate them to microstructure environments. Indeed, the MMD-MRI framework considers both *R*_1_ and *R*_2_ in the inversion and allows quantifying the components within the range of probed echo and repetition times. The limit of this quantification is dictated by the minimum τ_E_ which can induce an underestimation of water pools with short T_2_ leading to a low remaining signal at the minimum τ_E_ of 21 ms or an underestimation of water pools with short T_1_ leading to full recovery at the minimum τ_R_ of 1 second. The two GM components do not fall within those regimes and present only relatively small differences in relaxation rates and their relative quantification is expected to be meaningful. The *S*_0_ fraction of GM1 is thus approximately 3 to 10 times less intense than the one of GM2 with important disparities between brain regions. The GM1 fraction is especially intense in the cerebellum and olfactory bulb while GM2 is more present in the rest of the brain but shows a decrease in voxels close to the WM tracts. The superimposition of GM1 over GM2 shown in Supplementary Fig. 7 highlights this effect.

Even with the correlation of all those parameters, the attribution of the GM fractions to specific microstructures remains difficult especially since this case highlights a contradiction in the current literature. Indeed, in earlier OGSE studies, the diffusion-restriction was associated with intracellular diffusion in granule cells body due to the compatibility of the cell size with the restriction sensitivity range and their higher proportions in the brain regions highlighted by the restriction metrics such as the cerebellum^71,74,75^. This would lead to an assignment of GM2 as the intracellular water pool. However, other studies using multidimensional diffusion but without probing the diffusion-restriction were able to resolve multiple components in rat GM spinal cords^15,88^ and associated the slow diffusion component to intracellular water pools and the fast component to interstitial space. The direct translation of this conclusion to our case is contradictory with the OGSE-based assignment since *ω*-dependence is mainly observed in the fast (GM2) diffusion component. Note that this difference could also be attributed to the differences between the spinal cord and the rest of the brain. Other recent diffusion studies conducted with high *b*-values diffusion weighting^89,90^ also highlight a very low diffusivity compartment especially present in the gray matter human cerebellum with a proportion of up to 15 %^90,91^ which was attributed to cell bodies and thus intracellular water pools. Additional works demonstrate that such contrast can also be achieved with b-values up to 4 ms/µm^2^ only.^92^ A parallel with those results is appealing and would reinforce the hypothesis of the slow diffusion component being due to intracellular diffusion. However, the difference in subject, protocol, and fitted parameters prohibit a direct translation.

Further study remains necessary to attribute the two water pools to specific cell microstructures unequivocally. This could probably be achieved by either inducing cell swelling or sinkage^81^ or using an extracellular contrast agent^93^ to induce signal variations specific to microstructure and monitor them with the massively multidimensional diffusion relaxation correlation proposed here.

## Conclusion

Massively multidimensional diffusion MRI can be performed *in vivo* on rat brains with widely available preclinical MRI systems with moderate gradient strength. Its combination with state-of-the-art ultrafast EPI sequence allows multislice acquisition of a protocol of 389 images in only 17 minutes which can be translated to a high-resolution and low-distortion acquisition with segmented and full reversed phase-encode blips in an acquisition time slightly above two hours compatible with *in vivo* preclinical experiments. The use of modulated gradient shapes allows sampling a wide frequency range of 18 to 92 Hz at *b*-values up to 2.1 ms/µm^2^ and thus correlates the *ω*-dependent metrics usually obtained in OGSE with diffusion coefficient, shapes, orientation as well as T_1_ and T_2_ relaxation time. The wealth of information offered by this full correlation can be translated to numerous quantitative parameters maps proving a clear contrast between brain regions of different microstructures. The exploration of the multidimensional and multicomponent parameters distributions proving sub-voxel parameters quantifications allowed the identification of two separate components of GM with distinct diffusion and relaxation rate properties providing highly specific information for the study of healthy or diseased tissues.

## List of abbreviations

DTD: diffusion tensor distribution
D: diffusion tensor
*R*_1_: longitudinal relaxation rate
*R*_2_: transversal relaxation rate
τ_R_: repetition time
τ_E_: echo time
*ω*: diffusion frequency-dependence
b: diffusion encoding tensor
b(*ω*): diffusion encoding spectrum
*D*_iso_(*ω*): frequency-dependent isotropic diffusivity
*D*_Δ_(*ω*): normalized diffusion anisotropy
OGSE: oscillating gradient spin echo
MMD-MRI: massively multidimensional diffusion-relaxation correlation
SE-EPI: Spin Echo-Echo Planar Imaging
WM: white matter
GM: gray matter
CSF: cerebrospinal fluid
FLASH: Fast Low Angle shot
*b*_Δ_: encoding anisotropy
ob: olfactory bulb
lv: lateral ventricle
Cb: cerebellum
cc: corpus callosum
cg: cingulum
PAG: periaqueductal gray
ECIC: external cortex of the inferior colliculus
CBGr: cerebellar granule cell layer
CBML: cerebellar molecular layer

## Ethics

The research was conducted according to the principles expressed in the Declaration of Helsinki. The animal study was reviewed and approved by the Animal Committee of the Provincial Government of Southern Finland.

## Data and Code Availability

MATLAB source code for preprocessing and Monte-Carlo data inversion is freely available at https://github.com/maximeYon/MMD. The acquisition sequence is available upon reasonable request depending on Paravision versions.

## Author Contributions

MY: pulse sequence development, data acquisition and processing, manuscript drafting. ON: data acquisition and processing. DT: manuscript revision. AS: conceptualization, funding acquisition, project administration, supervision, manuscript revision. All authors contributed to the final version of the manuscript.

