## Supplementary information for "*In vivo* rat-brain mapping of multiple gray matter water populations using nonparametric D(*ω*)-*R*1-*R*2 distributions MRI"

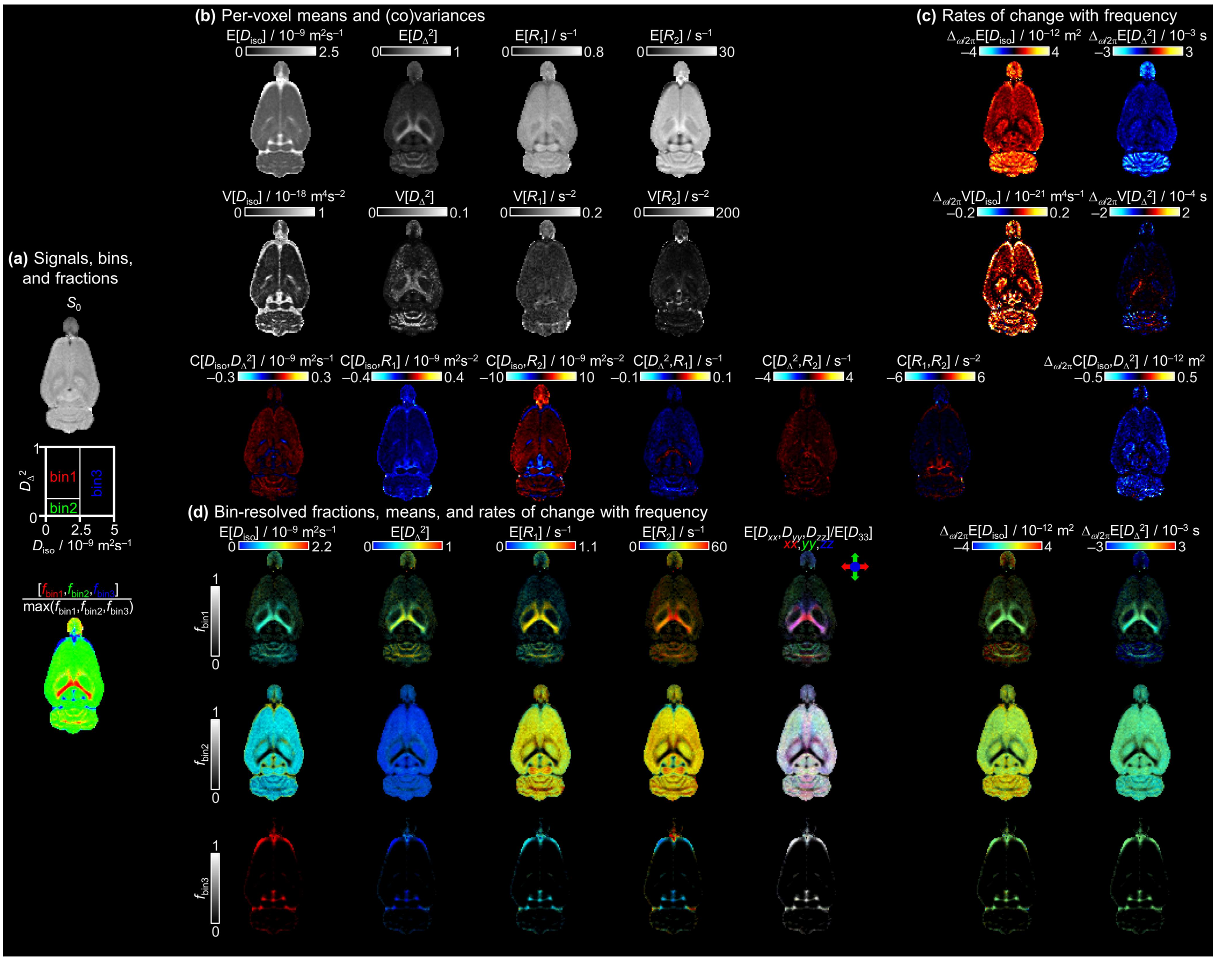


Supplementary Figure 1: Parameter maps derived from the per-voxel $\boldsymbol{D}(\omega)$-$R_1$-$R_2$-distributions of the in vivo rat brain slice 5. (a) Non-weighted signal $S_0 = S(b=0,\ \tau_R \rightarrow \infty,\ \tau_E = 0)$ obtained by Eq. 7; graphical representation of the 1/2/3 bins in the 2D $D_{iso}$-$D_\Delta^2$ space calculated according to the bin limit defined in the multidimensional diffusion data inversion section; and bin-resolved signal fractions $f_{bin1/bin2/bin3}$ coded into RGB color. The bin limits are selected to resolve white matter (WM), gray matter (GM), and cerebrospinal fluid (CSF). Primary colors indicate voxels containing pure WM, GM, or CSF while mixed colors show voxels with partial volumes of WM + GM (yellow), WM + CSF (purple), or GM + CSF (turquoise). (b) Per-voxel means E[x], variances V[x], and covariances C[x,y] of the $D_{iso}$, $D_\Delta^2$, $R_1$, and $R_2$ dimensions. (c) Per-voxel rates of change with frequency ($\Delta_{\omega/2\pi}$) means $\Delta_{\omega/2\pi}$E[x], variances $\Delta_{\omega/2\pi}$V[x], and covariances $\Delta_{\omega/2\pi}$C[x,y] of the $D_{iso}$, $D_\Delta^2$ dimensions. The ω-dependent parameters ($D_{iso}$ and $D_\Delta^2$) were calculated at the low sampling frequency of $\omega_{cent}$ = 18 Hz and their corresponding $\Delta_{\omega/2\pi}$ values between $\omega_{cent}$ = 18 and 92 Hz. (d) Bin-resolved signal fractions $f_{bin1/bin2/bin3}$, means (E[x] and $\Delta_{\omega/2\pi}$E[x]) coded into image brightness and color, respectively. The direction-encoded color is based on the lab-frame diagonal values $D_{xx}$, $D_{yy}$, and $D_{zz}$ normalized by the maximum eigenvalue $D_{33}$.

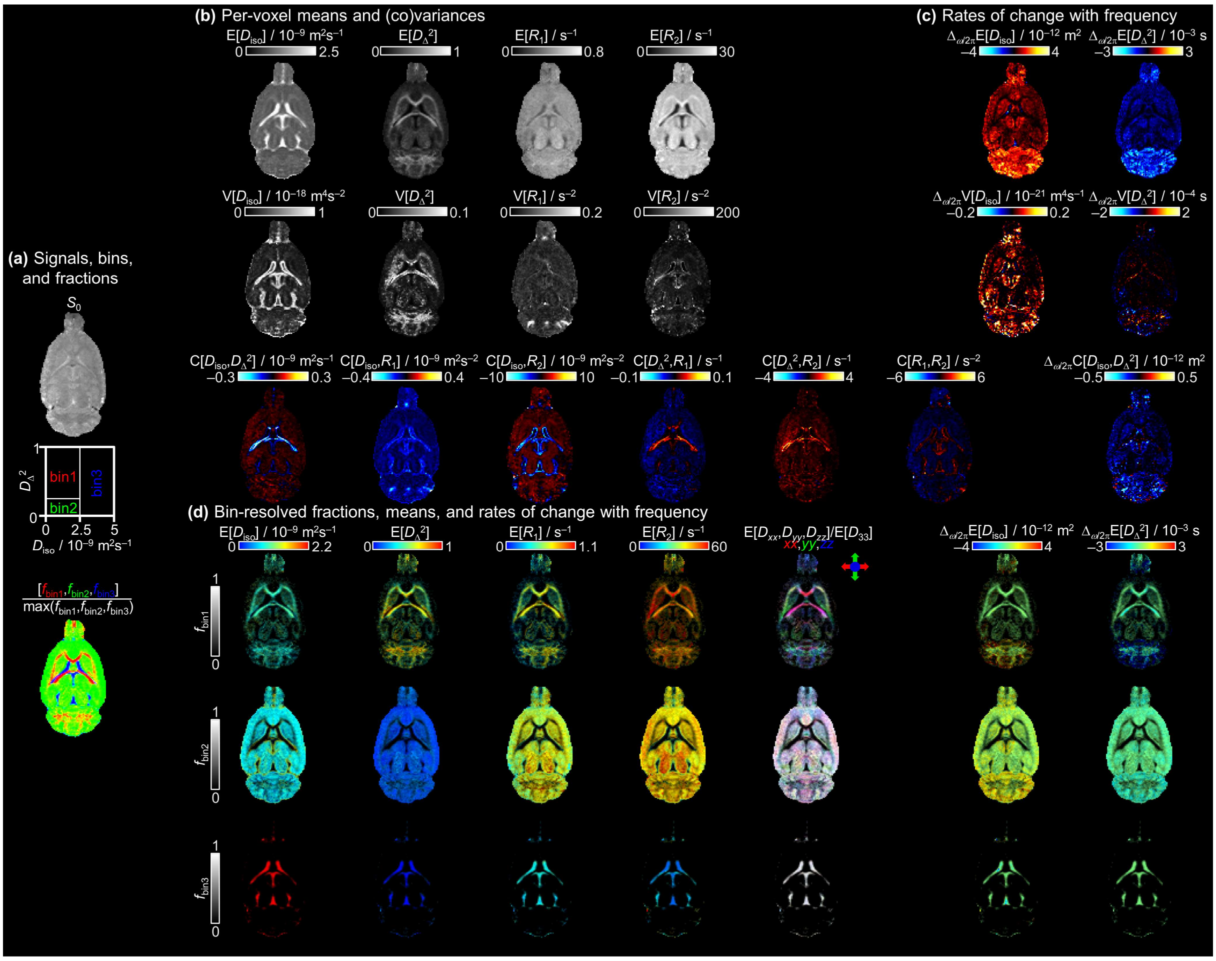

Supplementary Figure 2: Parameter maps derived from the per-voxel $\boldsymbol{D}(\omega)$-$R1$-$R2$-distributions of the in vivo rat brain slice 3.

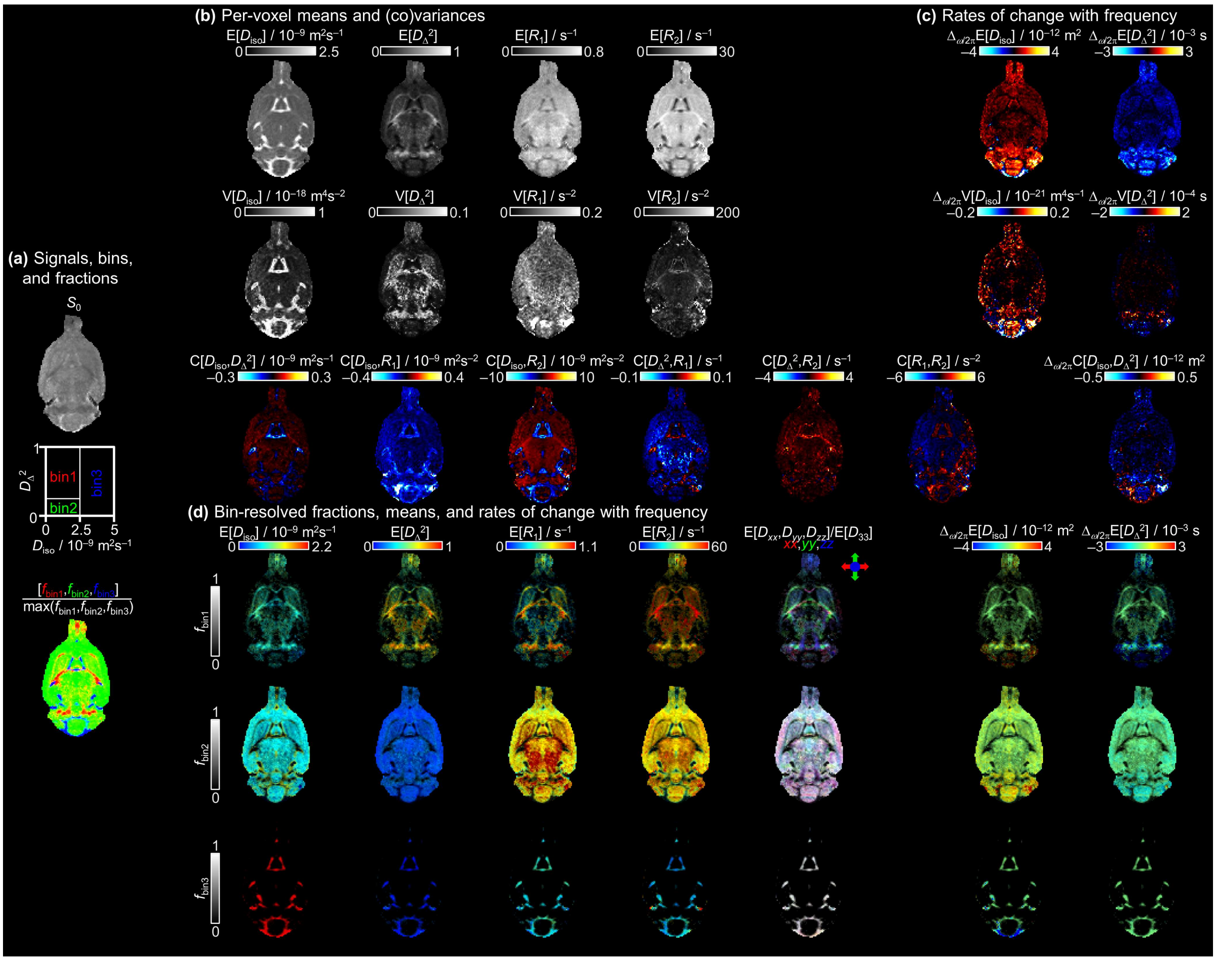

Supplementary Figure 3: Parameter maps derived from the per-voxel $\boldsymbol{D}(\omega)$-$R1$-$R2$-distributions of the in vivo rat brain slice 2.

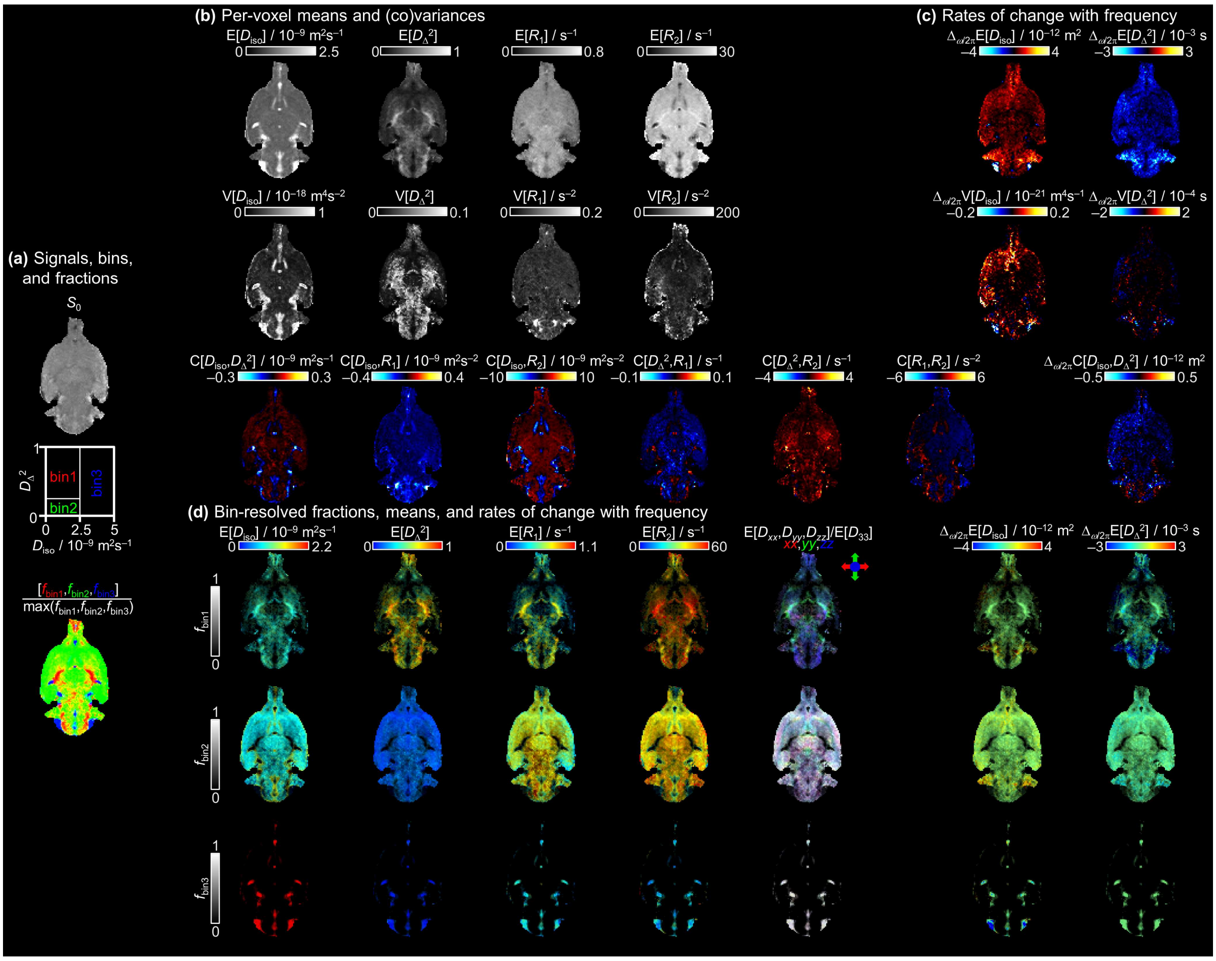


Supplementary Figure 4: Parameter maps derived from the per-voxel $\boldsymbol{D}(\omega)$-$R1$-$R2$-distributions of the in vivo rat brain slice 1.

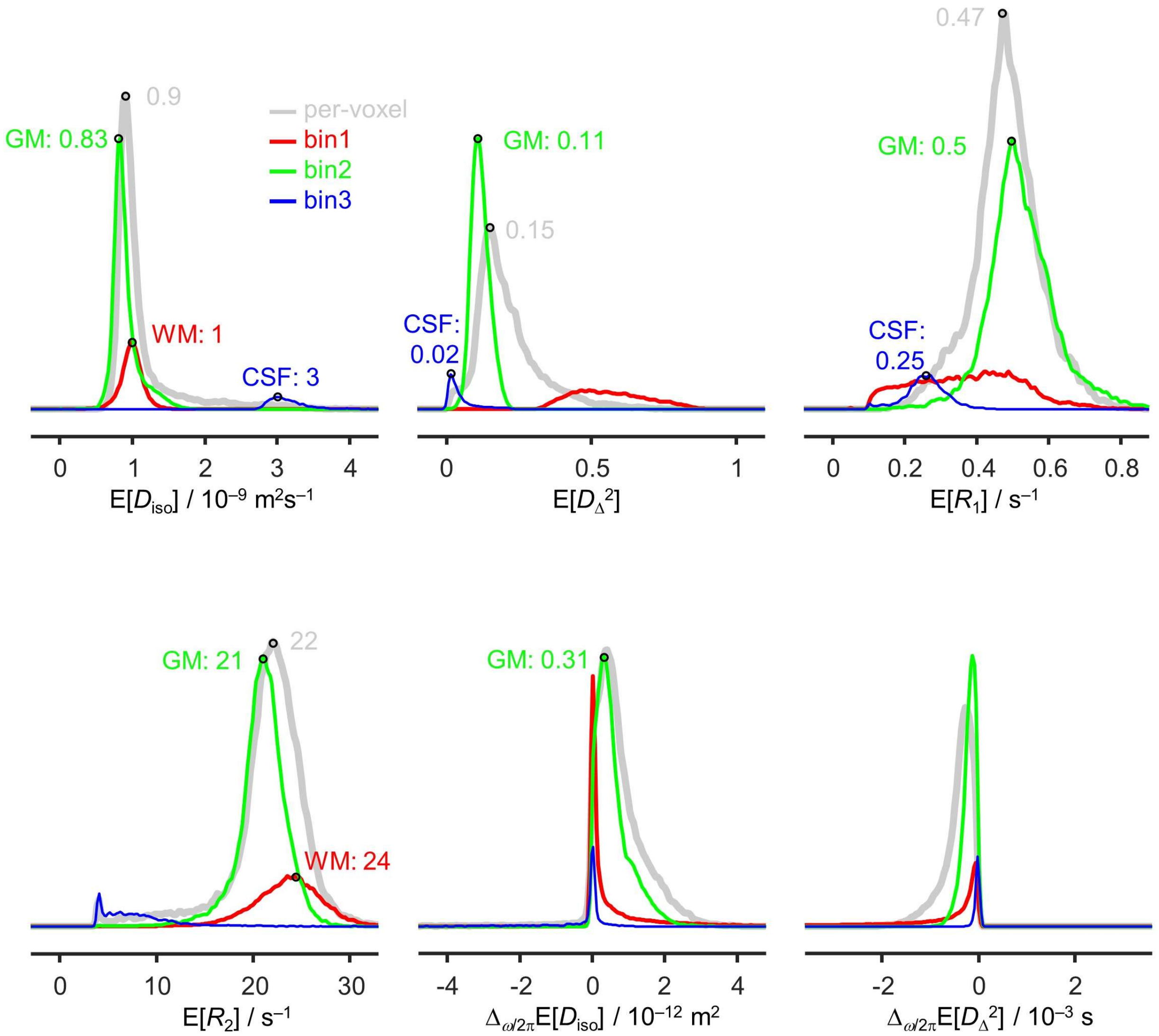


Supplementary Figure 5: Histograms of quantitative parameters median values across all voxels in the image. The histograms are obtained from the per-voxel (gray) and bin-resolved (red, green, and blue) maps of E[$D_{iso}$], E[$D_\Delta^2$], E[$R_1$], E[$R_2$], $\Delta_{\omega/2\pi}$E[$D_{iso}$], and $\Delta_{\omega/2\pi}$E[$D_\Delta^2$] in all slices and include weighting by $S_0$, $f_{bin1}$, $f_{bin2}$, and $f_{bin3}$. The abscissas cover the same ranges as the scale bars in Fig 3. Labeled points highlight representative values.

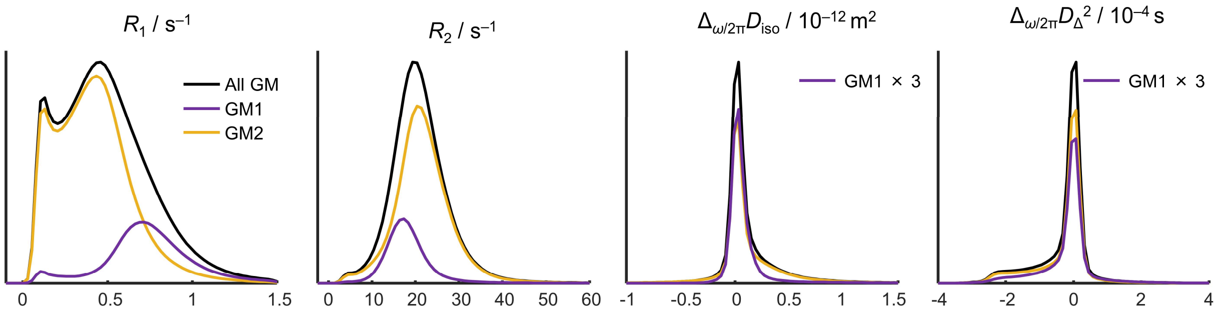


Supplementary Figure 6: $R_1$, $R_2$, $\Delta_{\omega/2\pi}D_{iso}$, and $\Delta_{\omega/2\pi}D_\Delta^2$ raw distributions of bin 2 in black, GM1 in purple, and GM2 in yellow in the 5 rat brain slices. In the two right plots, the intensity of the GM1 distribution was multiplied by three to ease the comparison of the low-intensity non-zeros points.

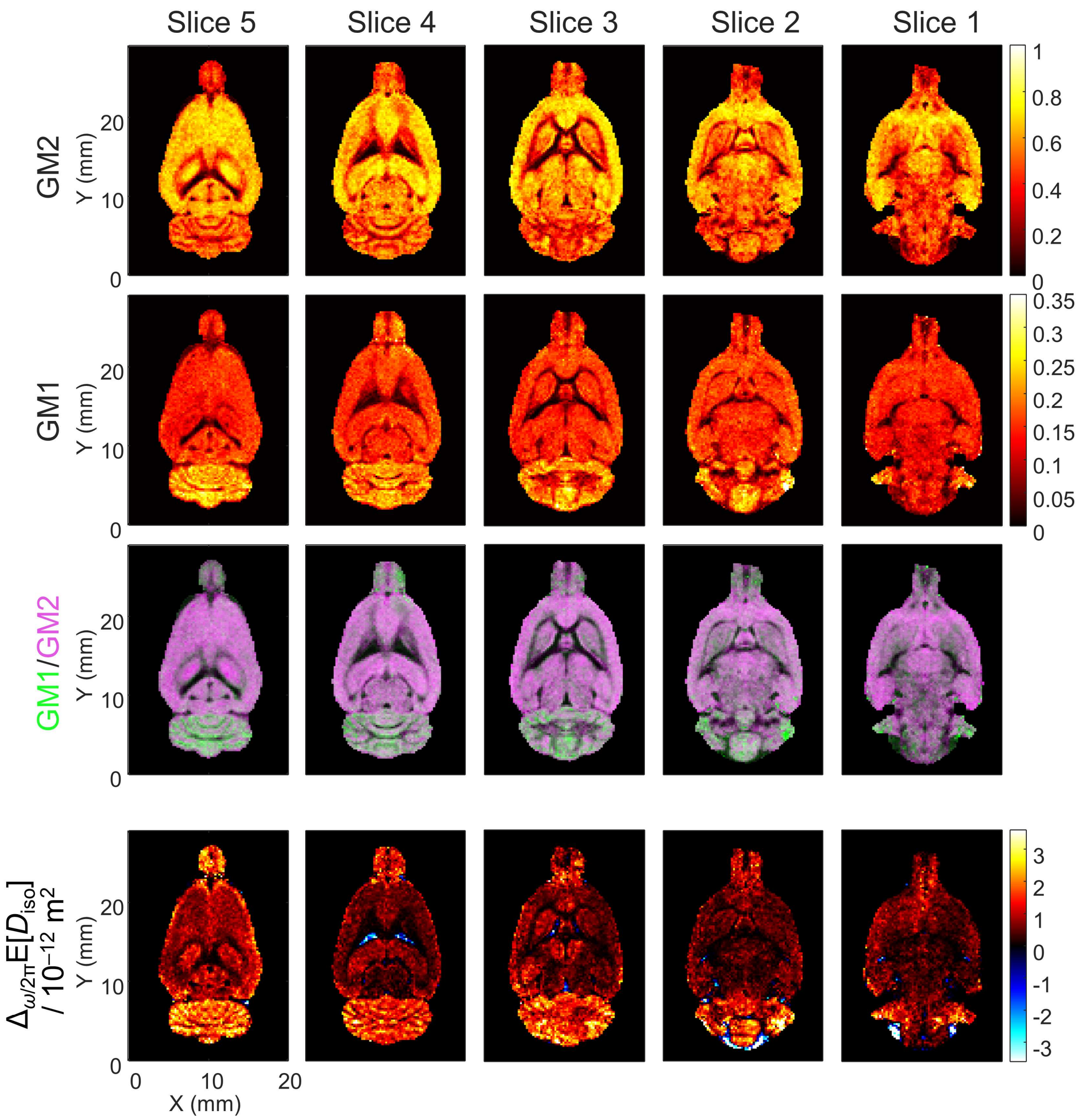


Supplementary Figure 7: parameter maps corresponding from top to bottom to: the $S_0$ fraction of GM2, the $S_0$ fraction of GM1, the overlap of GM1 over GM2 $S_0$ fractions normalized by their maximum intensities in green and purple respectively, and the $\Delta_{\omega/2\pi}E[D_{iso}]$ parameter maps reproduced here to ease comparison.
